# A virus associated with the zoonotic pathogen *Plasmodium knowlesi* causing human malaria is a member of a diverse and unclassified viral taxon

**DOI:** 10.1101/2024.09.18.613759

**Authors:** Mary E. Petrone, Justine Charon, Matthew J Grigg, Timothy William, Giri S Rajahram, Jacob Westaway, Kim A Piera, Mang Shi, Nicholas M. Anstey, Edward C. Holmes

## Abstract

Apicomplexa are single-celled eukaryotes that can infect humans and include the mosquito-borne parasite *Plasmodium*, the cause of malaria. Increasing rates of drug resistance in human-only *Plasmodium* species are reducing the efficacy of control efforts and antimalarial treatments. There are also rising cases of *P. knowlesi*, the only zoonotic *Plasmodium* species that causes severe disease and death in humans. Thus, there is a need to develop additional innovative strategies to combat malaria. Viruses that infect non-*Plasmodium* spp. disease-causing protozoa have been shown to affect pathogen life cycle and disease outcomes. However, only one virus (Matryoshka RNA virus 1) has been identified in *Plasmodium*, and none have been identified in zoonotic *Plasmodium* species. The rapid expansion of the known RNA virosphere using structure- and artificial intelligence-based methods suggests that this dearth is due to the divergent nature of RNA viruses that infect protozoa. We leveraged these newly uncovered data sets to explore the virome of human-infecting *Plasmodium* species collected in Sabah, east (Borneo) Malaysia. We identified a highly divergent RNA virus in two human-infecting *P. knowlesi* isolates that is related to the unclassified group ‘ormycoviruses’. By characterising fifteen additional ormycoviruses identified in the transcriptomes of arthropods we show that this group of viruses exhibits a complex ecology at the arthropod-mammal interface. Through the application of artificial intelligence methods, we then demonstrate that the ormycoviruses are part of a diverse and unclassified viral taxon. This is the first observation of an RNA virus in a zoonotic *Plasmodium* species. By linking small-scale experimental data to large-scale virus discovery advances, we characterise the diversity and genomic architecture of an unclassified viral taxon. This approach should be used to further explore the virome of disease-causing Apicomplexa and better understand how protozoa-infecting viruses may affect parasite fitness, pathobiology, and treatment outcomes.

## INTRODUCTION

Parasitic protozoa are a highly diverse collection of single-celled eukaryotes that can cause disease in many vertebrates. Organisms belonging to the phylum Apicomplexa are associated with a range of human diseases including malaria (*Plasmodium*), inflammation of the brain (*Toxoplasma*)^1^, diarrhea (*Cryptosporidium*)^2^, and severe anaemia (*Babesia*)^3^. *Plasmodium* is the leading cause of death from Apicomplexa in humans worldwide^4^. This mosquito-borne infection is estimated to have caused over 240 million cases of malaria and to have killed over 600,000 people in 2022 alone^5^.

Efforts to control and treat malaria are challenged by the complex ecology of this parasite^6^ and mounting antimalarial resistance^7,8^. Of the five human-only infecting species of *Plasmodium* (*P. falciparum*, *P. vivax*, *P. malariae*, *P. ovale wallikeri*, *and P. ovale curtisii*), *P. falciparum* and *P. vivax* cause the greatest morbidity and mortality, with *P. falciparum* accounting for more than 95% of malaria fatalities^4^. Partial resistance of *P. falciparum* to artemisinin is entrenched in the Greater Mekong subregion of Southeast Asia^9,10^ and has now emerged independently in Africa^5,11,8^. Eight additional *Plasmodium* species can cause human malaria through zoonotic transmission via mosquito vectors^12^. Among these, *P. knowlesi* is the only species to cause severe disease and death in humans^4,13–15^. Although predominating in Malaysian Borneo^16,17^, *P. knowlesi* is now recognised as a significant cause of malaria across Southeast Asia^18^, in association with changing land use and deforestation^18–21^, and in areas with declining incidence of the cross-protective species, *P. vivax*^22^. Thus, innovative strategies are needed to combat and control *Plasmodium* as the efficacy of accessible treatments declines in the human-only species, and changes in land-use cause greater numbers of zoonotic malaria cases.

One potential approach for malaria control involves the use of viruses that infect disease-causing protozoa. In a similar manner to how bacteriophage have been leveraged to combat drug-resistant bacterial infections^23–25^, protozoa-infecting viruses have been proposed as a potential new avenue for therapeutics^26,27^. These parasitic protozoan viruses (PPVs)^27^ have been identified in *Giardia*, *Leishmania*, *Cryptosporidum*^28,29^, *Eimeria*^30–38^, *Toxoplasma*^39^, *P. vivax*^40,41^, and *Babesia*^42,43^. They are of particular interest because some impact the parasite life cycle and modulate disease outcomes in the parasite host. Notably, *Leishmania* species that harbour Leishmania RNA virus 1 have been associated with an increased risk of treatment failure in humans^44^ and more severe disease outcomes in mice^45^. Similarly, it has been proposed that infection of *Toxoplasma* with the recently characterised apocryptoviruses (*Narnaviridae*) may be associated with increased disease severity in humans^39^, although this has yet to be formally tested. Cryptosporidium parvum virus 1 modulates the interferon response in *Cryptosporidium*-infected mammals^46^. To date, however, only one virus, Matryoshka RNA virus 1, has been identified in a *Plasmodium* species (*P. vivax*)^40,41^, and it is not known whether this virus impacts *Plasmodium* fitness or disease pathogenesis in humans.

Extending the known diversity of PPVs requires innovative approaches to virus discovery because both protozoa and the viruses that infect them are likely ancient and often highly divergent. As a case in point, the ormycoviruses were first identified in parasitic protozoa and fungi using structure-based methods^47^ and have since been identified in kelp (Stramenopila)^48^, ticks^49^, palm^50^, and additional fungal species^51,52^. This group of bi-segmented RNA viruses shares no measurable phylogenetic relationship to known viral taxa, rendering it invisible to sequence-based discovery methods^47^. Little else is known about ormycoviruses including their complete host range or whether they encode positive- or negative-sense genomes. The application of artificial intelligence-based methods^53^, in addition to large-scale sampling of aquatic environments^54^, has further uncovered previously inaccessible virus diversity, including entirely novel “supergroups” of unclassified viral taxa^53^. These tools and the data they have generated can be leveraged to explore the viromes of disease-causing protozoa including *Plasmodium*.

In this study, we combine these facets of virus discovery to characterise a divergent virus associated with human-infecting *P. knowlesi* isolates. We contextualise this virus within the vast viral diversity revealed through large-scale virus discovery studies. We also explore the complex ecology of viruses that infect parasites and can be transmitted as passengers to mammalian hosts. Our findings extend the diversity of known *Plasmodium*-associated viruses and highlight the importance of integrating large- and small-scale virus discovery research to better understand viruses that infect these ancient, microscopic hosts.

## RESULTS

### Identification of a divergent RNA virus associated with human-infecting *Plasmodium knowlesi*

To extend the known diversity of RNA viruses in disease-causing Apicomplexa, we analysed the metatranscriptomes of 18 human blood samples with PCR-confirmed *Plasmodium* infections and six uninfected human controls, collected in Sabah, east (Borneo) Malaysia between 2013 and 2014. These samples are the same as those previously described^40^. Of the patients with malaria, seven were infected with *P. vivax*, six with *P. knowlesi*, and five with *P. falciparum*^40^. Sequencing libraries were pooled according to *Plasmodium* species as were the negative controls, resulting in four libraries (SRR10448859-62, BioProject PRJNA589654). Matryoshka RNA virus 1 was previously found exclusively in all seven *P. vivax* isolates (SRR10448862)^40^.

We searched each library for divergent viruses using the RdRp-scan bioinformatic pipeline^55^. This revealed a putative, highly divergent RNA-dependent RNA polymerase (RdRp) that was 3,177nt in length with a complete open reading frame (ORF) and robust sequencing coverage in the *P. knowlesi* library (SRR10448860) (**Fig. 1a**). No identical or related sequences were found in the other three libraries. The transcript was relatively abundant (1.4% of non-rRNA reads), and we confirmed the presence of this putative RdRp in two of the six isolates in the pool using RT-PCR (**Table S1, Fig. S1**). Both patients with putative virus-infected *P. knowlesi* isolates were from Kota Marudu district residing in villages approximately 30km apart. There was three months difference in the date of hospital presentation. Both had uncomplicated malaria with parasitemia of 7,177 and 41,882 parasites/µL, respectively, which were higher than the median parasitemia found in the *P. knowlesi* infections that lacked the putative virus (4,518/µL). Parasitemia was correlated with the RdRp signals we observed with PCR (**Fig. S1**).

**Figure 1.**
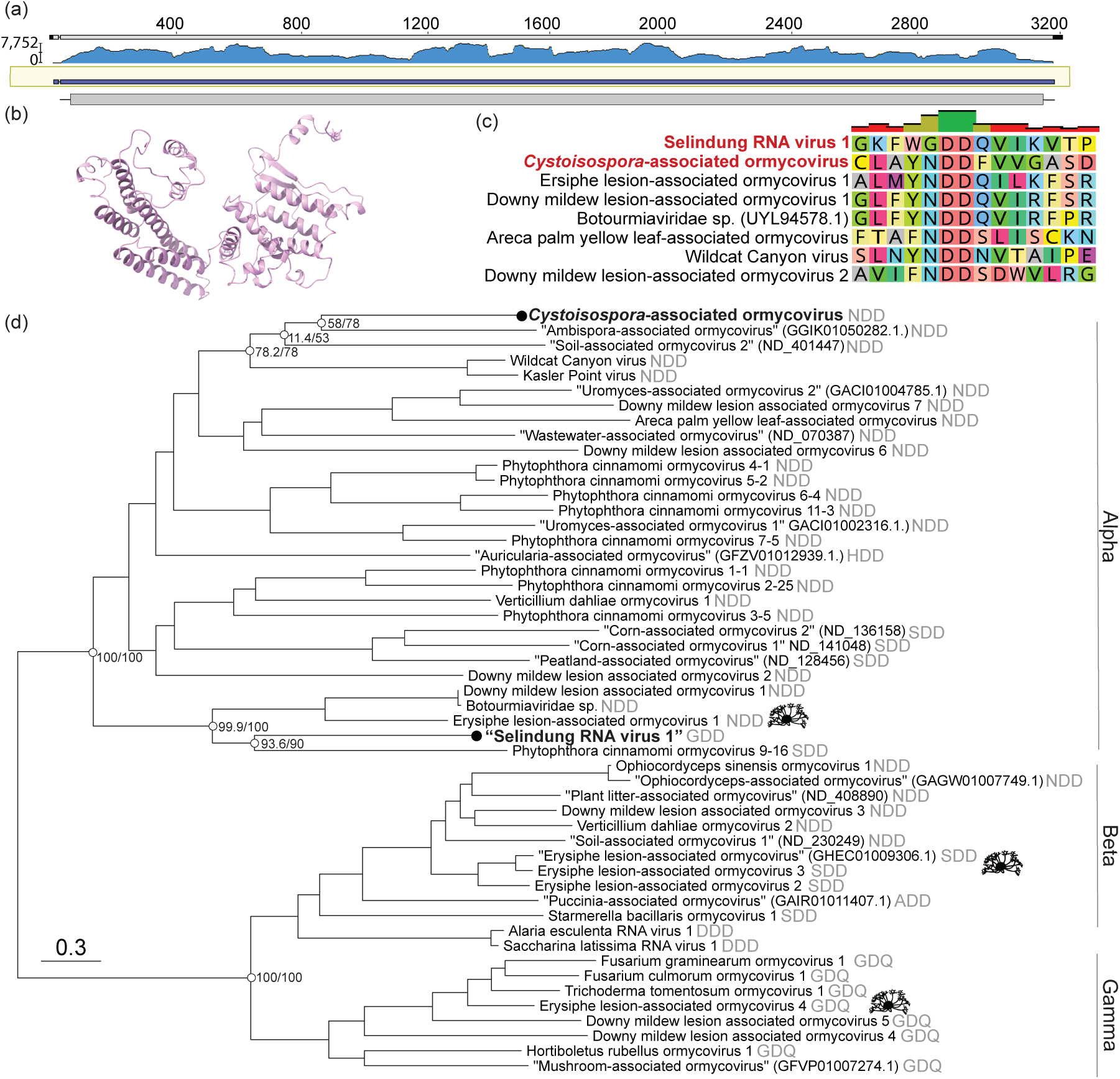
A divergent RNA virus associated with human-infecting *P. knowlesi* is a member of the unclassified group ‘ormycovirus’. (a) Sequencing coverage of the RNA-dependent RNA polymerase (RdRp) of a *P. knowlesi*-associated viral contig. Trimmed reads were mapped to the assembled contig using BBMap^63^ and visualised with Geneious Prime v2024.0.7. (b) The predicted structure of the putative hypothetical protein of Selindung RNA virus 1. (c) MAFFT alignment of motif C in the palm domain of the *P. knowlesi*- and *Cystoisospora*-associated viruses and representative ormycoviruses. (d) Phylogenetic inference of the ormycoviruses aligned with MAFFT. The positions of *Erysiphe*-associated viruses are denoted with black icons (source: phylopic.org). Black tip dots indicate viruses identified in this study. Tips with names in quotes were previously identified but not named^47^. Their corresponding NCBI or RVMT accession is shown in parentheses. The catalytic triad encoded in each palm domain is denoted in grey. Support values are shown at select nodes as sh-aLRT/UFBoot. Tree branches are scaled to amino acid substitutions.

Further inspection indicated that this putative virus was a bi-segmented ormycovirus likely infecting the *Plasmodium*. The divergent RdRp shared low but detectable similarity with that of seven previously identified viruses, of which six were ormycoviruses (**Table S2**). We identified a putative second segment of unknown function, 1,721nt in length sharing 22.8% identity (e-value = 3.14 × 10^−15^) with the hypothetical protein of Erysiphe lesion-associated ormycovirus 1 (USW07196). The structures of the putative and known hypothetical proteins were significantly similar (p-value = 1.62 × 10^−2^) when predicted with AlphaFold2^56,57^ and compared by pairwise alignment with FATCAT^58^ (**Fig. 1b**). Similar transcripts were not identified in the ormycovirus-negative libraries from the same BioProject. Analysis of the library composition with CCMetagen^59^ and the KMA database^60^ did not reveal plausible host candidates aside from the *Plasmodium*, which comprised 24% of non-rRNA reads. The remainder aligned to the *Hominidae*, reflecting that the *Plasmodium* were themselves infecting humans. We assumed that the host range of the ormycoviruses likely did not extend to vertebrates, consistent with their absence in the humans without *Plasmodium* infection. Unlike its closest relatives, the *P. knowlesi*-associated RdRp encoded GDD in motif C of its palm domain rather than NDD (**Fig. 1c**).

To assess the prevalence of this and other ormycoviruses in *P. knowlesi*, we screened 1,470 *P. knowlesi* RNA SRA libraries (**Supp. Data 1**) with a custom ormycovirus database. This returned no additional ormycovirus candidates. However, all 1,470 libraries were generated from only seven BioProjects, and only the library we generated was derived from human-host *P. knowlesi* infections. The majority (n = 1,356) were generated from macaque-host *P. knowlesi* infections, and all of these were generated by a single contributor from a small set of laboratory-maintained Rhesus macaques (PRJNA508940, PRJNA526495, and PRJNA524357). Sixty-one libraries were derived from cell culture, and the source of 52 (BioProject PRJEB24220) could not be determined. Thus, an accurate prevalence estimate of the *P. knowlesi*-associated ormycovirus could not be obtained from this data set.

We next investigated the prevalence of ormycoviruses more broadly in disease-causing Apicomplexa by screening 2,898 RNA SRA libraries (*Cryptosporidum*, *Coccidia*, *Toxoplasma*, *Babesia*, and *Theileria*) (**Supp. Data 2**). This yielded identical ormyco-like RdRp segments in the transcriptomes of 22 Coccidia (*Cystoisospora suis*) libraries, 21 of which belonged to the same BioProject (PRJEB52768)^61^. The remaining library (SRR4213142) was published by the same authors, suggesting that all 22 libraries were generated from the same source^62^. The transcripts of the *Cystoisospora*-associated virus encoded complete ORFs with an NDD motif C and were ∼3.1kb in length (range: 3009-3203) (**Table S3**). This virus was highly divergent, sharing only 32.5% identity (e-value = 6 × 10^−37^) with its closest blast hit (Wildcat Canyon virus, WZL61396.1). It was also at low abundance across the 22 libraries (range: 0.01-0.08% of non-rRNA reads). We could not conclude that *C. suis* was the host because fungi represented 4.6% of the non-rRNA reads in a representative library (ERR9846867). Regardless, the prevalence of ormycoviruses was 100% among *Cystoisospora suis* libraries but otherwise very low in this data set (0.76%).

Phylogenetic analysis placed both Apicomplexa-associated viruses in the “Alpha” clade of the ormycoviruses (**Fig. 1d**). The topology of the inferred phylogenies was stable across six combinations of alignment and trimming methods and recapitulated the three main ormycovirus clades “Alpha”, “Beta”, and “Gamma”^47^ with strong support (**Fig. S2**). Viruses did not cluster by host. For example, viruses associated with the fungal species *Erysiphe* fell across all three clades and encoded three different catalytic triads (**Fig. 1d, *icons***), and the Apicomplexa-associated viruses were not closely related within the Alpha clade.

We concluded that the *P. knowlesi*-associated virus represents the first evidence of an RNA virus associated with *P. knowlesi* and constitutes only the second instance of an RNA virus associated with any *Plasmodium* species. We have provisionally named it “Selindung RNA virus 1” because it appeared to be concealed (“terselindung”, Bahasa Malaysia) within the *Plasmodium* parasite, and we will use this name herein.

### Ormycoviruses are associated with arthropod metatranscriptomes

In addition to expanding the diversity of *Plasmodium*-associated RNA viruses, Selindung RNA virus 1 was of particular interest because it had evidently been transmitted along with its *Plasmodium* host to a human via a mosquito vector. Taking this together with the detectable phylogenetic relationship of this virus and two viruses recovered from tick metagenomes (Wildcat Canyon virus and Kasler Point virus), we posited that ormycoviruses might exhibit a complex ecology at the arthropod-mammal interface. We therefore sought to further extend the known host range of ormycoviruses to the transcriptomes of the arthropods that indirectly transmit them.

We screened the 4,864 arthropod libraries available on NCBI Transcriptome Shotgun Assemblies (TSA) as of August 2024, initially using Kasler Point virus (WZL61394) as input and then following an iterative process (see *Methods*). In this way we identified 15 putative viruses associated with three of the four extant subphyla of the Arthropoda: Chelicerata (n = 1), Crustacea (n = 1), and Hexapoda (n = 13) (**Table S4**). All shared detectable but minimal sequence similarity with published ormycoviruses (range: 27.1-41.0%, **Table S4**). Two encoded GDD at motif C like Selindung RNA virus 1, while the remainder had NDD at this position.

Phylogenetic analysis again supported the conclusion that these viruses are part of the ormycovirus group (**Fig. 2a**). All viruses identified in this study fell in the Alpha clade. Selindung RNA virus 1 formed a group with the other two GDD-encoding viruses (Beetle-associated ormycovirus 1 and Bristletail-associated ormycovirus 1). This placement was consistent across all six iterations of phylogenetic inference (**Fig. S3**). However, aside from this instance and the Gamma clade (GDQ), minimal clustering of motifs was observed. In addition, although the host organisms had been collected from all six inhabited continents, there was no clustering of viruses by geographic region of sampling (**Fig. 2a**).

**Figure 2.**
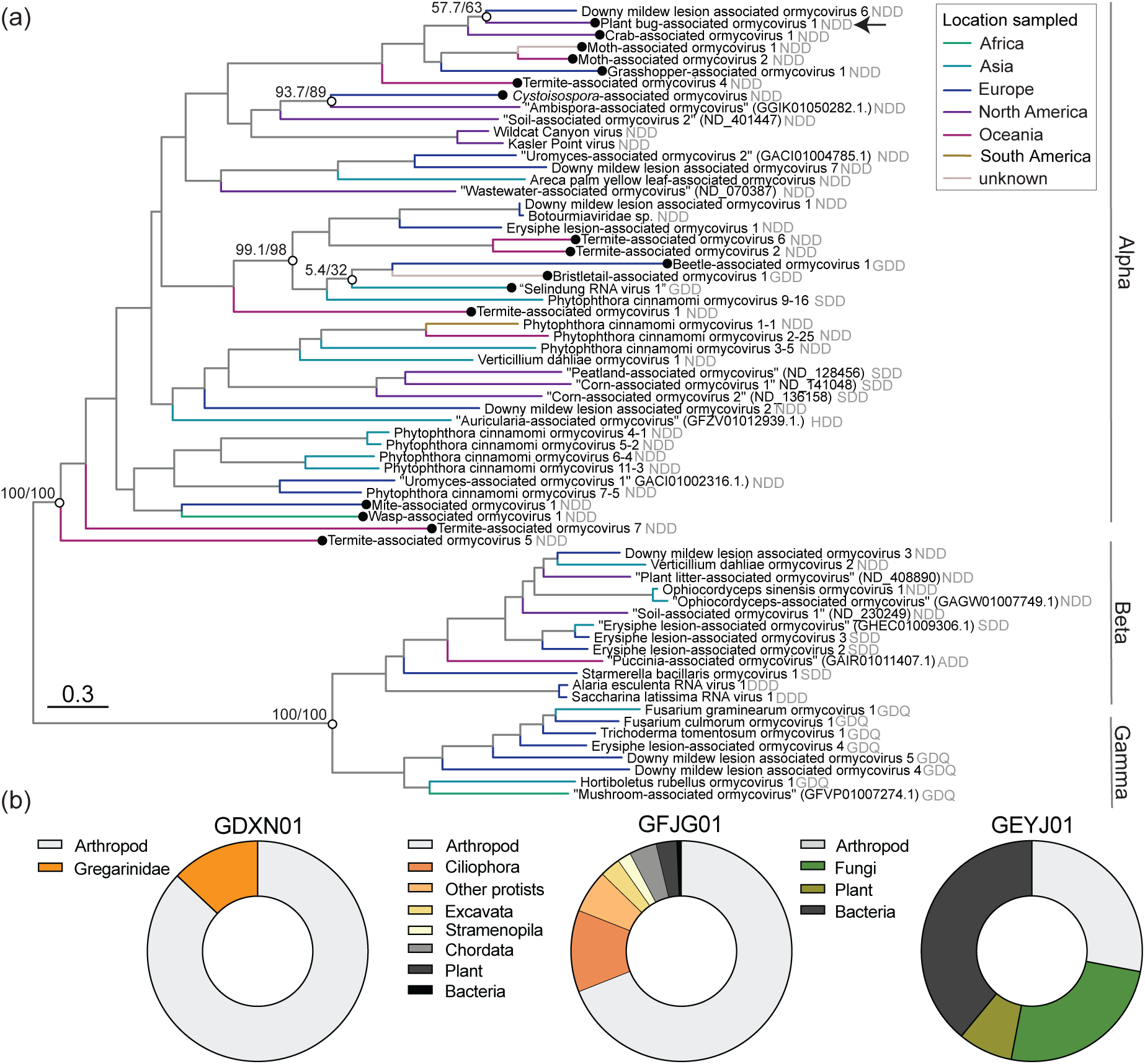
Ormycoviruses are associated with arthropod transcriptomes. (a) Phylogenetic inference of the extended diversity of ormycoviruses. Viruses identified in this study are indicated by black circles. The arrow indicates the position of the virus that appears to use the ciliate genetic code. Clades are annotated according to designations established by Forgia *et al*.^47^. The catalytic triad encoded in each palm domain is denoted in grey. The tip labelling scheme for unnamed viruses (denoted by quotation marks) is the same as in Fig. 1. Support values are shown at select nodes as sh-aLRT/UFBoot. Tree branches are coloured by the location where each tip was sampled, and they are scaled by amino acid substitutions. (b) Library composition of select arthropod assemblies. The graph labels correspond to the TSA project ID.

We concluded that at least some of these viruses were likely infecting single-celled organisms rather than the arthropods themselves for two reasons. First, assessment of each library composition revealed instances of parasitic hosts. Contigs mapping to alveolates accounted for more than one tenth of one Hexapoda (GDXN01) and the only crustacean (GFJG01) library (13% *Gregarinidae* and 12% Ciliophora, respectively) (**Fig 2b**). Similarly, the mite assembly (GEYJ01) included 25% of contigs mapping to fungi (**Fig. 2b**). Second, the virus identified in GBHO01 (*Lygus hesperus*) likely utilised the ciliate genetic code (i.e., only a truncated ORF could be recovered with the standard genetic code) yet fell within the diversity of the taxon (**Fig. 2a, Fig. S3, *arrow***). Identical amino acid translations of the crustacean-associated virus were produced when either the standard or the ciliate genetic code were used.

As with the *P. knowlesi* library, we searched these assemblies for hypothetical proteins. From this, we identified a putative second segment in the *Machilis pallida* (Hexapoda) assembly HBDP01 containing Bristletail-associated ormycovirus 1 that was 1,619bp in length and encoded a partial ORF (HBDP01002991.1). We could not recover candidates corresponding to the remaining libraries or assemblies.

### The ormycoviruses are members of a diverse and unclassified viral taxon

The wide host range of the ormycoviruses, spanning Alveolata, Stramenopila, and Opisthokonta (Fungi), suggested that this unclassified group harboured unrealised viral diversity. We therefore aimed to contextualise the diversity of the ormycoviruses within unclassified taxa identified in virus discovery studies. To do this, we assembled a custom database of the viruses identified using an artificial intelligence-based method^53^ and screened the ormycoviruses against it using DIAMOND Blastx^64^. This approach placed ormycoviruses within an unclassified taxon referred to in the original study as the proposed “SuperGroup 024”^53^, a name which we will use herein.

Phylogenetic analysis illustrated that the current set of ormycoviruses represent only a fraction of the total diversity of this group as they fell throughout the phylogeny. Interestingly, the addition of the SuperGroup 024 viruses expanded the diversity of the Alpha group, scattering the original members across three sections of the tree (**Fig. 3a, *blue branches***). The Beta and Gamma clades were unchanged and characterised by a long branch at their shared base (**Fig. 3a, *green and yellow branches***).

**Figure 3.**
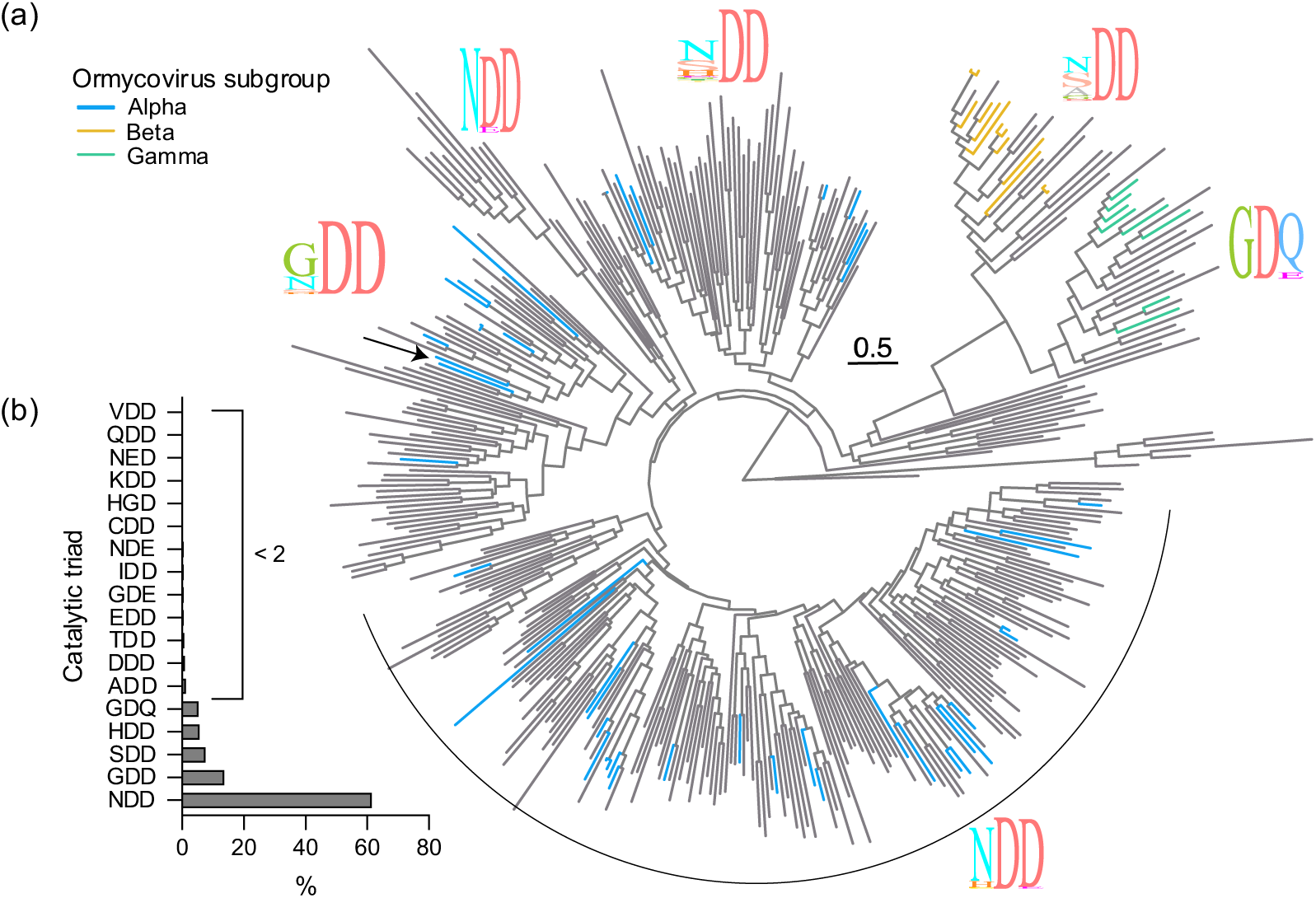
Ormycoviruses are members of a diverse and unclassified viral taxon with a flexible motif C in its palm domain. (a) Phylogenetic inference of viruses in SuperGroup 024^53^. Branches are coloured by their placement in the ormycovirus-only phylogenetic tree (Fig. 1 and 2). Grey tree branches indicate that those tips were not previously recognised as ormycoviruses. The icons show the proportion of individual amino acids at each position of the catalytic triad in motif C of the RdRp palm domain for the corresponding clades. The arrow indicates the topological position of Selindung RNA virus 1. Tree branches are scaled according to amino acid substitutions. (b) Distribution of catalytic triads encoded by members of SuperGroup 024. The x-axis shows the percentage that each triad comprises among all known SuperGroup 024 species.

Members of SuperGroup 024 encoded a more diverse set of catalytic triads at the motif C palm domain compared to the original ormycovirus data set^47^ (**Fig 3b**). However, their addition did not lead to observable clustering of discrete motif sequences, as flexibility was observed throughout the phylogeny. Selindung RNA virus 1 again fell in a section predominated by GDD at that position (**Fig. 3a, *arrow***).

We searched the libraries containing SuperGroup 024 RdRp segments for ormycovirus hypothetical proteins. Of the 259 SRA libraries in which SuperGroup 024 RdRps were detected and assembled, we recovered hypothetical protein candidates at least 1000bp in length in 190 (73.4%). It was not possible to assign hypothetical proteins to corresponding RdRps as many libraries contained multiple RdRp segments. Despite this, our finding supports the conclusion that bisegmentation is a characteristic of viruses in this taxon and that ormycoviruses and SuperGroup 024 are one and the same.

## DISCUSSION

This study expands the diversity of *Plasmodium*-associated RNA viruses and presents the first evidence of an RNA virus associated with zoonotic transmission of *P. knowlesi*. Previously, only Matryoshka RNA virus 1 (*Narnaviridae*) had been identified in a *Plasmodium* species (the human-only *Plasmodium* species, *P. vivax*)^40,41^. Although it is not possible to conclusively establish that Selindung RNA virus 1 was infecting the parasite from metatranscriptomic data alone, lines of indirect evidence suggest that it was. Most notably, no other probable hosts, including fungi, were identified in the library, and the RdRp contig was relatively abundant (1.4% of non-rRNA reads). Contamination was an unlikely source because neither the putative RdRp nor the second segment were detected in the other three libraries extracted and sequenced at the same time. In addition, we were able to confirm the presence of the RdRp segment in two of the six *P. knowlesi* isolates using PCR. We therefore concluded that Selindung RNA virus 1 most likely represents an RNA virus in a second *Plasmodium* species.

Robust sampling of natural *P. knowlesi* infections is needed to evaluate the prevalence and pathobiology of Selindung RNA virus 1. We observed one instance of Selindung RNA virus 1 among 1,470 SRA libraries, which suggests that associations occur infrequently and contrasts with the identification of Matryoshka RNA virus 1 in 13 of 30 *P. vivax* SRA libraries^40^. However, ours was the only library to have been generated from isolates collected from naturally infected humans, while most of the publicly available data were derived from laboratory experiments. The detection of the RdRp segment in two of six isolates in our library could indicate that associations are more frequent in natural infections in Sabah, but our study was not powered to assess this. Similarly, whether the observation that presence of the virus was correlated with higher parasitemia is meaningful requires further epidemiological investigation.

Arthropods are a powerful tool for measuring the prevalence of viruses in nature, particularly when sampling from humans or other vertebrates is not feasible. The identification of ormycoviruses in arthropod metatranscriptomes and in a human blood sample suggests that these viruses represent a unique type of arbovirus that can be transmitted as a passenger between arthropods and mammals. Mosquito-based surveillance methods have been proposed for tracking the incidence and spread of human pathogens^65,66^. Unlike cell culture or primary samples, which rely on symptomatic individuals with access to diagnostic testing, arthropod-based surveillance would be relatively unbiased, enabling more accurate estimates of protozoan virus prevalence and diversity within communities. When combined with cell culture data, this approach could also be used to parse arthropod- and protozoan-infecting viruses. Because they can be indirectly transmitted by arthropods, it may be that other protozoan viruses have already been identified, but their relationship to their protozoan host was obscured because they were part of an arthropod metatranscriptome.

An incidental and surprising finding was the identification of an ormycovirus that appears to use a non-standard genetic code (Plant bug-associated ormycovirus 1). Despite this difference, the virus fell within the diversity of the ormycoviruses and SuperGroup 024. As RNA viruses are reliant on host machinery for translation, it was previously proposed that the evolution of alternative genetic codes was an antiviral defence^67^. Under this assumption, the use of host-specific genetic codes by RNA viruses would imply a long-term virus-host coevolutionary relationship, and we would not expect to find viral taxa in which members use different genetic codes. Genetic code switching has been observed infrequently in the *Picornavirales* and *Lenarviricota*^68^. Whether these select instances are an aberration in an otherwise broadly held rule of virology requires further investigation. However, we posit that there may be many more instances of code switching within known viral taxa that have been overlooked as a consequence of inadequate bioinformatic workflows. For example, if we had used an automated pipeline that filtered out contigs that did not produce an ORF with the standard genetic code, Plant bug-associated ormycovirus 1 would have been removed from our data set. We therefore advocate for the inclusion of multiple genetic codes when searching for divergent RNA viruses.

That Selindung RNA virus 1 does not belong to a known viral taxon is notable because it demonstrates that parasitic protozoa likely harbour currently unrealised diversity, and additional discoveries may be imminent as new bioinformatic tools are developed to explore the RNA virosphere. However, the discovery of the ormycoviruses highlights the importance of linking large-scale metatranscriptomic data to smaller-scale experimental work when searching for protozoan viruses. Large-scale virus discovery studies often prioritise environmental samples such as water^54^, sediment^53^, and soil^68^ because these biodiverse sources are rich with RNA viruses. Yet, this approach cannot distinguish between bacterial-, archaeal-, and eukaryotic-infecting RNA viruses. Without the discovery of the ormycoviruses and the experimental validation by Forgia *et al*.^47^, SuperGroup 024 would have been overlooked as a potential source of protozoan virus candidates. Similarly, large-scale studies are not equipped to distinguish segmented from non-segmented viruses because they necessarily focus on detecting RdRps, rendering them “blind” to segmentation. The molecular characterisation of the ormycoviruses again demonstrates this limitation because their hypothetical protein does not share detectable sequence or structural similarity with known viral proteins. Without the incidental finding by Forgia *et al*.^47^, we would not have been able to infer that the ormycoviruses and the members of SuperGroup 024 are likely segmented viruses.

This, as with other metagenomic studies, primarily serves to generate hypotheses and raise questions about RNA virus evolution and biology that require additional experimental data to answer. It is not known whether the ormycoviruses are positive- or negative-sense viruses. Forgia *et al*. proposed that they are negative-sense because they observed a higher proportion of negative-sense RNA in their samples^47^; however, this is not definitive. The presence of both SDD and GDD catalytic triads in motif C in the palm domain counters the hypothesis that SDD is specific to segmented negative-sense RNA viruses^69^, although it is possible that ormycoviruses do indeed fall into this category. The flexibility of the catalytic triad also raises the question of whether individual triads have a detectable impact on the biology of the virus and why flexibility is permitted in otherwise highly conserved region of the virus genome. From a global health perspective, the most important questions to address include how viral infection of *Plasmodium* affects onward *Plasmodium* transmission and the pathobiology of *Plasmodium* in humans. Additionally, which part of the parasite the virus infects and whether this could be used as a potential drug target remain unanswered. It has already been shown that viruses can serve as a weapon against drug-resistant bacterial infections^23–25^. Whether a similar approach could be deployed to combat malaria and other disease-causing Apicomplexa should be a research priority.

## METHODS

### Human malaria isolates

*Plasmodium* RNA was isolated from cryopreserved red cells collected from 18 patients with acute malaria, enrolled in Kudat Division, Sabah, Malaysia in 2013 and 2014^15^. PCR was used to confirm *Plasmodium* species as *P. knowlesi* (n=6), *P. vivax* (n=7) and *P. falciparum* (n=5), as previously reported^40^.

### SRA library data sets

#### BioProject PRJNA589654 libraries

*Plasmodium* SRA libraries in BioProject PRJNA589654 (n = 4) (i.e., the BioProject that contained Matryoshka RNA virus 1) were downloaded from NCBI. Nextera adapters were trimmed using Cutadapt v.1.8.3^70^ with the parameters removing 5 bases from the beginning and end of each read, a quality cutoff of 24, and a minimum length threshold of 25. The quality of trimming was assessed using FastQC v0.11.8^71^. rRNA reads were removed using SortMeRNA v4.3.3^72^, and non-rRNA reads were assembled using MEGAHIT v1.2.9^73^.

#### Disease-causing Apicomplexa libraries

We downloaded all *P. knowlesi* RNA SRA libraries of at least 0.5Gb in size available on NCBI as of August 2024 (n = 1,470). We also downloaded all RNA SRA libraries for *Cryptosporidium, Coccidia, Toxoplasmosis, Babesia,* and *Theileria* available on NCBI as of March 2024 that are at least 0.5Gb in size and generated on the Illumina platform (n = 3,162).

#### SuperGroup 024 libraries

To analyse the libraries containing RdRp segments of so-called SuperGroup 024^53^, we first downloaded all of the contigs designated in this group by Hou *et al*.^53^ (http://47.93.21.181/). We then extracted the corresponding SRA libraries from each sequence header and removed duplicates (n = 273). All but one were downloaded from NCBI. The library SRR1027962 failed repeated attempts to download, likely due to its size (99.8Gb).

### Arthropod Transcriptome Shotgun Assemblies (TSA) screen

We began by screening all arthropod TSA (n = 4,864) available in August 2024, using Kasler Point virus (a tick-associated ormycovirus) as input. This screen was performed with tBLASTn implemented in the NCBI Blast web interface (https://blast.ncbi.nlm.nih.gov/Blast.cgi). All hits were reviewed and filtered according to three criteria: (1) the contig was at least 800bp in length, (2) the contig encoded an uninterrupted ORF, (3) the contig did not return any hits to cellular genes when screened against the NCBI non-redundant (nr) database. We then aligned our filtered data set using MAFFT^74^ with default parameters, and selected the most divergent virus according to the distance matrix. This virus was then used as input for an additional screen of the arthropod TSA. This process was repeated until no new contigs were identified.

### Library processing

#### Contig assembly

For all data sets obtained from the SRA, Nextera adapters were trimmed using Cutadapt v.1.8.3^70^ with the parameters described above. The efficacy of trimming was assessed using FastQC v0.11.8^71^. In total, 1,470 *P. knowlesi* libraries, 2,898 additional Apicomplexa libraries, and 259 libraries included by Hou *et al.*,^53^ were successfully assembled using MEGAHIT v1.2.9^73^.

#### Abundance estimates

The expected count of putative viral transcripts was inferred using RSEM v1.3.0^75^. For the *P. knowlesi* library containing the ormycovirus (SRR10448860), reverse-strandedness was specified to match the sequencing protocol. Default parameters were used for the remaining libraries. To infer the proportion of reads of each putative viral transcript, we calculated the total expected count for the isoforms in each library and used this value as the denominator to measure the percentage that putative viral reads comprised in the library. This analysis was performed in R v4.4.0.

### Identification of divergent viruses

#### Polymerase segment identification

We identified Selindung RNA virus 1 using the RdRp-scan workflow^55^. Briefly, we screened the protein sequence and HMM-profile of assembled contigs from each library against a viral RdRp database. To search for additional divergent viruses, we screened all SRA libraries against the RdRp-scan database^55^ and a custom database containing known ormycoviruses using DIAMOND Blastx v2.0.9^64^ and the setting ‘ultra-sensitive’. This database included the 39 published ormycoviruses and the Selindung RNA virus 1 RdRp segment. Only hits with e-values below 1e-07 were retained for further analysis. Contigs with hits to this database were then screened against the NCBI nr protein database to remove false positives, again using DIAMOND Blastx v2.0.9^64^ and an e-value threshold of 1e-07. The parameter ‘very-sensitive’ was specified. Contigs that shared detectable sequence similarity to cellular genes were excluded from further analysis. Nucleotide sequences were translated using Expasy (https://web.expasy.org/translate/). The standard genetic code was used by default. Contigs that did not return an ORF in any frame with this code were checked manually using all codes available in Expasy.

#### Second segment identification

We first used blastn to screen libraries for contigs sharing conserved 5’ and 3’ termini of the corresponding ormycovirus RdRp. When this did not reveal any candidates, we compiled a database of all known ormycovirus second segments and used this to screen all SRA libraries using DIAMOND Blastx v2.0.9^64^. Contigs that had statistically significant hits to this database were checked against the NCBI nr protein database to remove false positives (i.e., cellular genes). Nucleotide sequences were either translated individually with Expasy (https://web.expasy.org/translate/) or with InterProScan v5.65-97.0. For sequences processed with the latter, the longest translated ORFs were used for downstream analysis. To tally the number of SuperGroup 024 libraries with detectable hypothetical proteins, we cross-checked the presence of RdRp segments and hypothetical protein segments in each library using R v4.4.0.

For the primary *P. knowlesi* library, we searched for similar sequences to those at the 5’ and 3’ termini of the RdRp segment in other contigs in the library. To do this, we extracted these regions from the RdRp segment and used each as input for tblastn against the assembled library (SRR10448860). To ensure that the putative Selindung RNA virus 1 hypothetical protein was not present in other libraries in the same BioProject, we used this sequence as input for tblastn against the three remaining libraries.

Both tblastn screens were implemented in Geneious Prime v2024.0.7 and default parameters were used.

#### PCR validation

We first generated cDNA from the isolates using the SuperScript IV reverse transcriptase (Invitrogen). These products were then used as templates for amplification with PCR. Reactions were carried out in a total volume of 50ul, of which 25ul was SuperFi II (Invitrogen) master mix and 1ul was the cDNA template. 2.5ul of forward and reverse primers were used (**Table S1**). Reactions were performed on a thermocycler with the following conditions: 98°C for 1 min followed by 35 cycles of 98°C for 10s, 60°C for 10s, 72°C for 1 min, and 72°C for 5 min. The PCR products were analysed on an agarose gel. We used *Plasmodium* LDHP primers as the positive control.

### Library composition analysis

#### CCMetagen

The composition of individual sequencing libraries was assessed using ccmetagen v1.2.4^59^ and kma v1.3.9a^60^ using assembled contigs as input. The results presented in **Fig. 2b** were visualised with Prism v.10.3.0.

### Protein structure inference

The structure of the putative hypothetical proteins of Selindung RNA virus 1 and Erysiphe lesion-associated ormycovirus 1 were predicted using AlphaFold2^56,57^ implemented in the Google Colab cloud computing platform. The confidence (as measured by pIDDT) of the prediction was compared across five models, and the highest performing models (Selindung RNA virus 1: #2, Erysiphe lesion-associated ormycovirus 1: #4) were selected for downstream analysis (**Fig. S4**). To assess structural similarity, we performed a pairwise alignment of the resulting pdb files of each predicted structure using FatCat^58^. All pdb files were visualised in ChimeraX v1.7.1^76^.

### Functional domain inference

Several approaches were used to infer functional domains in the hypothetical protein, although none were successful. We first performed a preliminary check with InterProScan^77^, screening against the CDD, NCBIfam, and TMHMM databases. This approach was implemented in Geneious Prime v2024.0.7. We then employed Phyre2^78^ and HHPred^79^ using PDB. Finally, we used the predicted structure of the hypothetical protein of Selindung RNA virus 1 as input for FoldSeek^80^, implemented on the Foldseek Server.

### Phylogenetic analysis

To assess the phylogenetic relationships of the ormycoviruses identified in this study with those documented previously, we compiled a data set of all known ormycoviruses. This comprised 36 ormycoviruses^47–52^ and unclassified or misclassified ormycoviruses that shared detectable sequence similarity with known ormycoviruses: Wildcat Canyon virus (WZL61396), Kasler Point virus (WZL61394), and a fungus-associated “Botourmiaviridae” (UYL94578). For the SuperGroup 024 analysis, we utilised the data set featured in the phylogenetic analysis presented by Hou *et al.*^53^.

We first added the *P. knowlesi*- and *Cystoisospora-*associated viruses identified in this study to the ormycovirus data set and aligned with MAFFT v7.490^81^ and MUSCLE v5.1^82^. Ambiguities in each alignment were considered in three ways using trimAl v1.4.1^83^: (i) no ambiguities were removed; (ii) ambiguities were removed using a gap threshold of 0.5 and a conservation percentage of 50; (iii) ambiguities were removed using the parameter “gappyout”. Phylogenetic trees for these six alignments were inferred using ModelFinder and IQ-TREE v1.6.12^84^. To quantify support for the topology, we again used 1000 ultra-fast bootstraps and 1000 SH-aLRT bootstrap replicates.

To infer the pan-SuperGroup 024 phylogeny, all amino acid sequences were aligned using both MAFFT v7.490^81^ and MUSCLE v5.1^82^. Ambiguities were removed using trimAl v1.4.1^83^ and the parameter -gappyout. The phylogenetic tree was inferred using IQ-TREE v1.6.12^84^ with ModelFinder limited to LG. Support values were measured with 1000 ultra-fast bootstraps (UFboot) and 1000 sh-aLRT bootstrap replicates.

All trees were visualised with ggtree^85,86^ (implemented in R v4.4.0) and Adobe Illustrator v26.4.1.

### Motif C tally and visualisation

The catalytic triad encoded by each virus in SuperGroup 024 was recorded and tabulated using R v4.4.0. The results were visualised with Prism v.10.3.0.

Sequences from individual clades were extracted from the SuperGroup 024 phylogeny by selecting individual nodes using the function “extract.clade()” implemented in the R package ape. Sequences from each clade were then realigned with MAFFT and the motif C logos were generated according to the consensus sequence in Geneious Prime v2024.0.7.

## DATA AVAILABILITY

All sequencing data analysed in this study are publicly available on NCBI (ormycoviruses) and an independent repository (http://47.93.21.181/, SuperGroup 024). Assembled contigs for the viruses identified in this study, the custom database used to screen libraries, alignments, and tree files are available on GitHub (https://github.com/mary-petrone/Plasmodium_ormyco).

## ACKNOWLEDGMENTS

This work was funded by a National Health and Medical Research Council (NHMRC) Investigator award (MJG), AIR@InnoHK administered by the Innovation and Technology Commission, Hong Kong Special Administrative Region, China (ECH), a Sydney ID Seed Funding Award (MEP), and the National Institutes of Health, USA R01 AI160457-01 and Malaysia Ministry of Health Grant BP00500/117/1002 (GSR). We thank the Director General of Health, Malaysia for the permission to publish this article.

We also thank Jon Mifsud for his BatchArtemisSRAMiner pipeline (https://github.com/JonathonMifsud/BatchArtemisSRAMiner), which we used for all SRA screens, and Dr. Alvin Kuo Jing Teo for suggesting the name Selindung RNA virus 1.

## AUTHOR CONTRIBUTIONS

M.E.P., J.C., N.M.A., and E.C.H. designed the study. M.J.G, T.W., G.R., J.W., and K.A.P. designed the original malaria studies and collected the samples. M.E.P., J.C., and M.S. performed the experiments and analyses. M.E.P. wrote the initial manuscript draft. All authors reviewed and edited the manuscript.

