## Supplementary material for "A virus associated with the zoonotic pathogen *Plasmodium knowlesi* causing human malaria is a member of a diverse and unclassified viral taxon": Fig S1

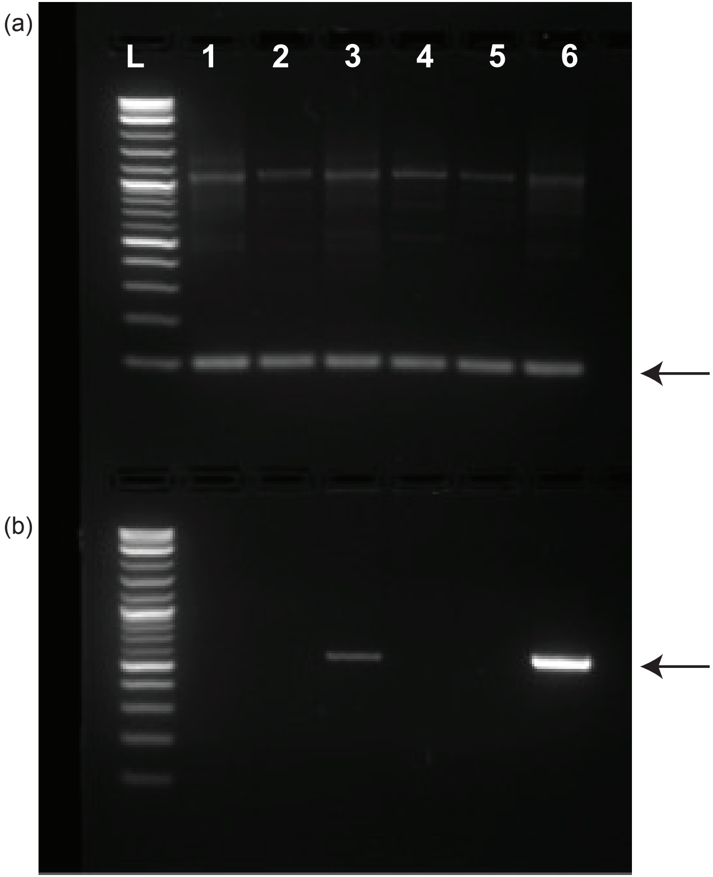


**Figure S1 Detection of divergent *P. knowlesi*-associated RNA virus in two *P. knowlesi* isolates collected from human blood samples.** (a) House-keeping gene *Plasmodium* LDHP (arrow). (b) Presence of the divergent RdRp segment in two *P. knowlesi* isolates (arrow).
