## Supplementary material for "A virus associated with the zoonotic pathogen *Plasmodium knowlesi* causing human malaria is a member of a diverse and unclassified viral taxon": Fig S2

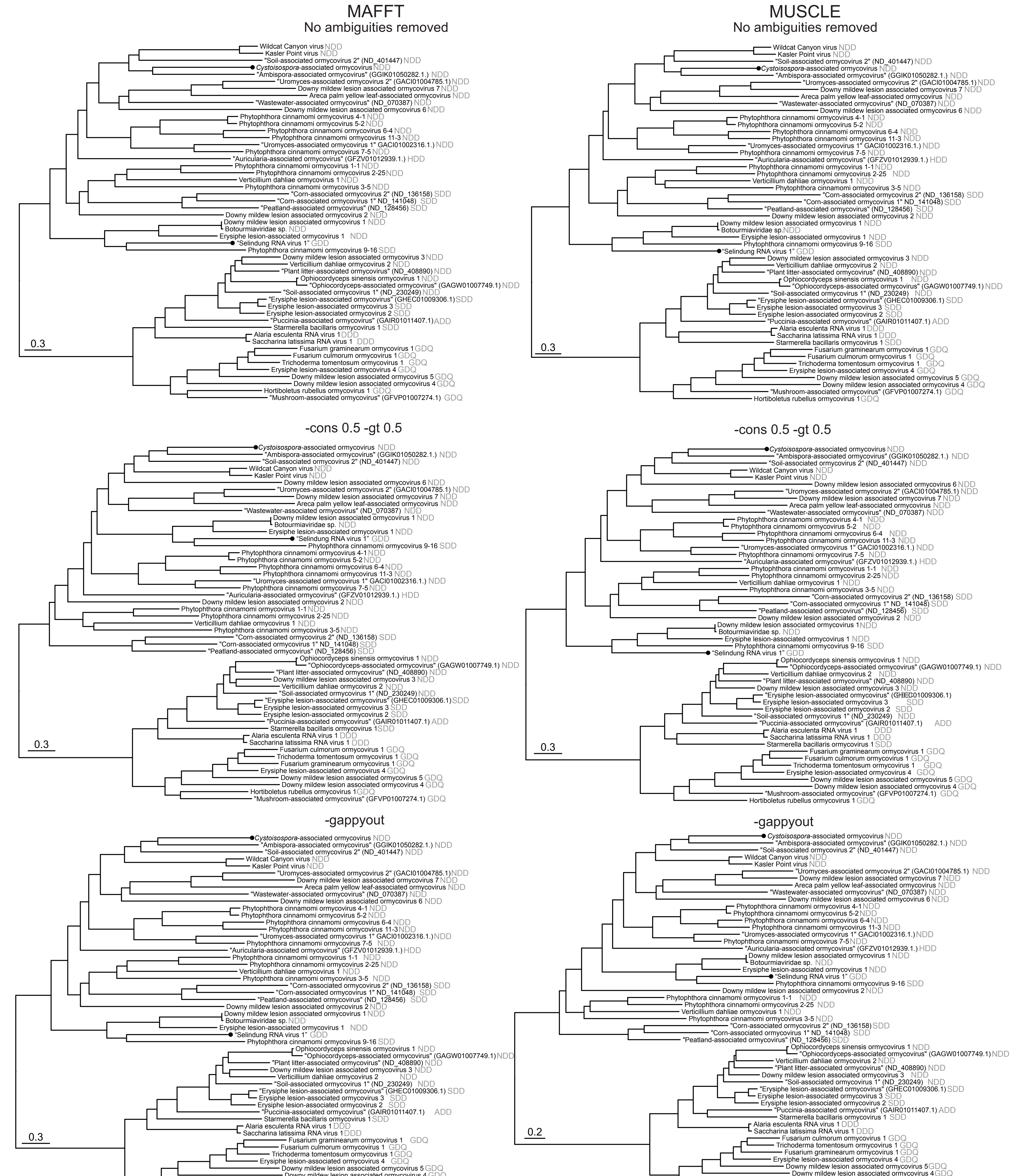


**Figure S2 *Plasmodium*- and *Cystoisospera*-associated ormyco-like viruses fall within the ormycoviruses across 6 combinations of aligning and trimming methods.** The Plasmodium-associated virus is denoted as "Selingdung RNA virus 1". Tree branches are scaled by number of amino acid substitutions and coloured by geographic region of sampling. Viruses identified in this study are denoted by black tips. Catalytic triads of the palm domain Motif C are shown in grey.
