## Supplementary material for "A virus associated with the zoonotic pathogen *Plasmodium knowlesi* causing human malaria is a member of a diverse and unclassified viral taxon": Fig S3

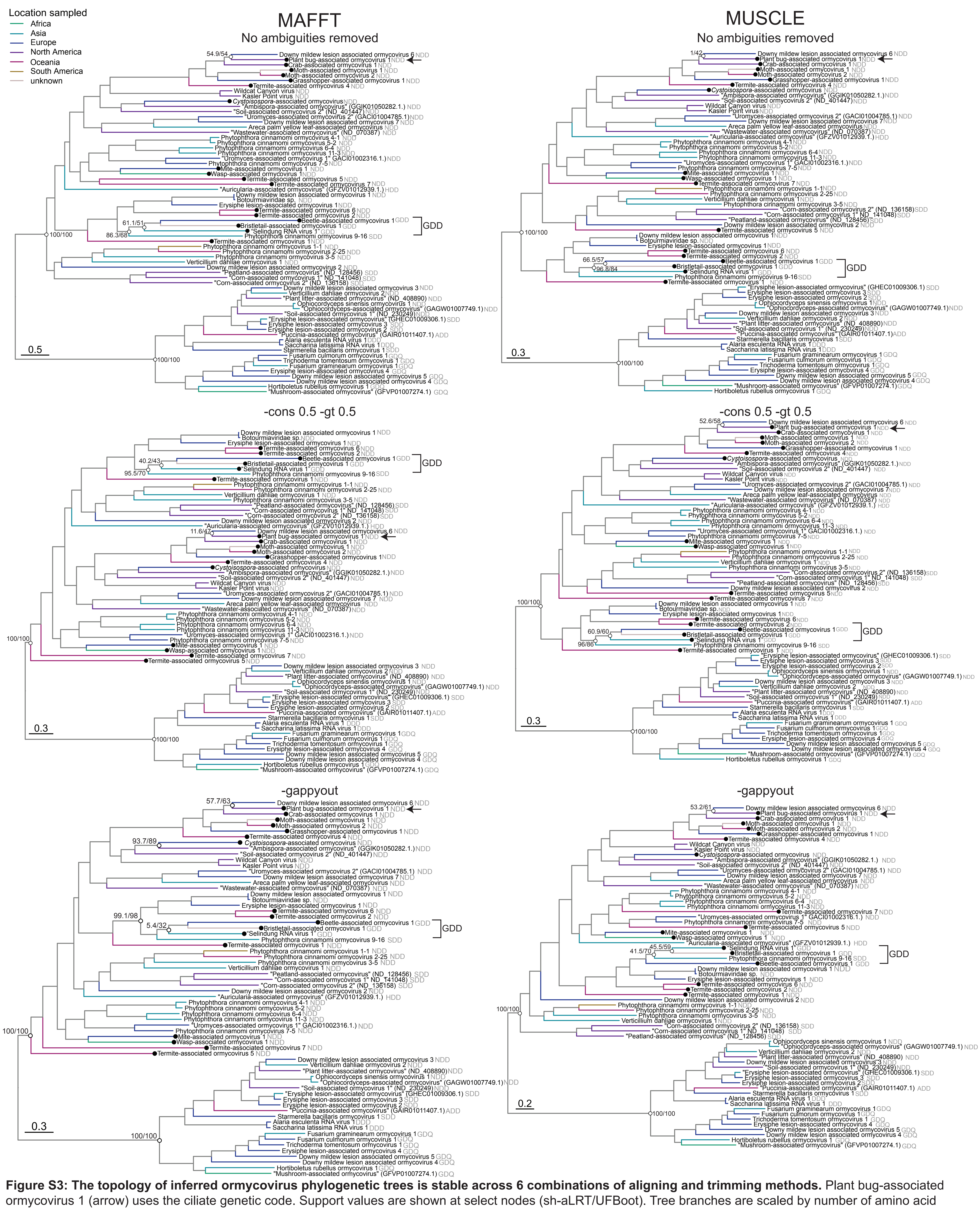


**Figure S3: The topology of inferred ormycovirus phylogenetic trees is stable across 6 combinations of aligning and trimming methods.** Plant bug-associated ormycovirus 1 (arrow) uses the ciliate genetic code. Support values are shown at select nodes (sh-aLRT/UFBoot). Tree branches are scaled by number of amino acid substitutions and coloured by geographic region of sampling. Viruses that encode GDD at their C motif domain are indicated with a bracket. Viruses identified in this study are denoted by black tips.
