## Supplementary material for "A virus associated with the zoonotic pathogen *Plasmodium knowlesi* causing human malaria is a member of a diverse and unclassified viral taxon": Fig S4

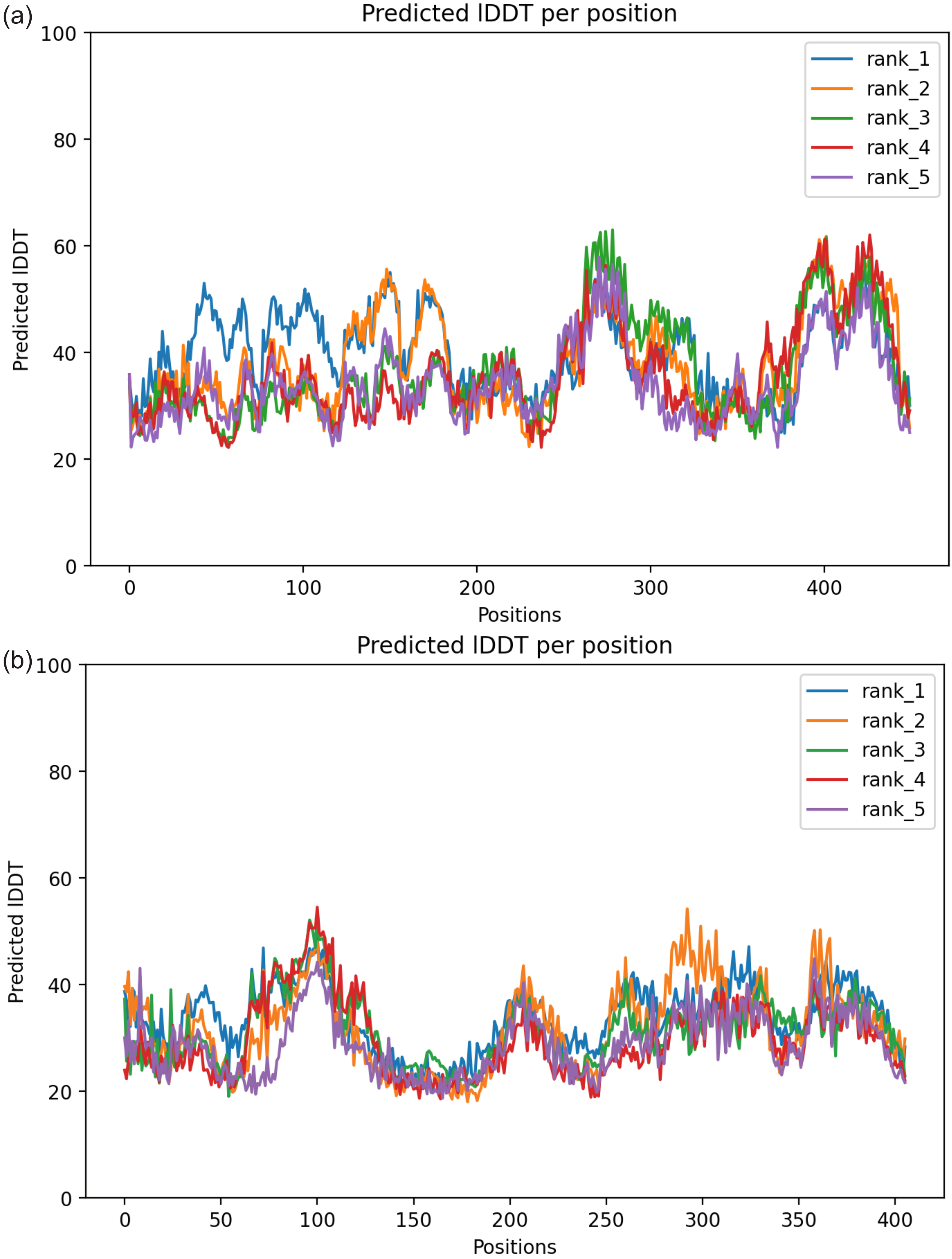


**Figure S4. Predicted Local Distance Difference Tests (lDDT) for 5 models of structural predictions of the hypothetical protein of the *P. knowlesi*-associated virus (a) and Erysiphe lesion-associated ormycovirus 1 (b) generated using ColabFold.**
