## Supplementary material for "A virus associated with the zoonotic pathogen *Plasmodium knowlesi* causing human malaria is a member of a diverse and unclassified viral taxon": Table S1

**Table S1 Primers targeting the *P. knowlesi*-associated ormycovirus used for RT-PCR**

| Primer direction | Primer sequence |
|---|---|
| Forward | CCTGGCTTTGGGGGCAATA |
| Reverse | CCCATCCCTCTGGAGTCCA |
