## Supplementary material for "A virus associated with the zoonotic pathogen *Plasmodium knowlesi* causing human malaria is a member of a diverse and unclassified viral taxon": Table S2

### Table S2 Blastx hits to P. knowelsi-associated virus

| Hit name | Query cover (%) | Identity (%) | E-value |
|---|---|---|---|
| Downy mildew lesion associated ormycovirus 1 (RdRp) | 49 | 31.78 | 3e-52 |
| Botourmiaviridae sp. (RdRp) (UYL94578.1) | 49 | 31.84 | 7e-52 |
| Erysiphe lesion-associated ormycovirus 1 (RdRp) | 49 | 32.70 | 4e-48 |
| Areca palm yello leaf-associated ormycovirus (RdRp) | 51 | 21.40 | 2e-04 |
| Downy mildew lesion associated ormycovirus 2 (RdRp) | 41 | 24.89 | 0.001 |
| Wildcat Canyon virus (RdRp) | 27 | 23.59 | 0.003 |
