## Supplementary material for "A virus associated with the zoonotic pathogen *Plasmodium knowlesi* causing human malaria is a member of a diverse and unclassified viral taxon": Table S3

**Table S3 Ormyco-like RdRp segments identified in 22 *Cystoisospora suis* libraries**

| Library | Segment Length |
|---|---|
| ERR9846864 | 3182 |
| ERR9846865 | 3120 |
| ERR9846866 | 3160 |
| ERR9846868 | 3176 |
| ERR9846869 | 3202 |
| ERR9846870 | 3145 |
| ERR9846872 | 3186 |
| ERR9846873 | 3186 |
| ERR9846874 | 3177 |
| ERR9846875 | 3175 |
| ERR9846876 | 3181 |
| ERR9846877 | 3203 |
| ERR9846878 | 3166 |
| ERR9846879 | 3175 |
| ERR9846880 | 3180 |
| ERR9846881 | 3181 |
| ERR9846882 | 3177 |
| ERR9846883 | 3161 |
| ERR9846884 | 3173 |
| SRR4213142 | 3185 |
| ERR9846867 | 3090 |
| ERR9846871 | 3182 |
