## Supplementary material for "A virus associated with the zoonotic pathogen *Plasmodium knowlesi* causing human malaria is a member of a diverse and unclassified viral taxon": Table S4

### Table S4 Summary of ormycoviruses identified in Arthropoda TSA libraries

| Contig ID | Assigned name | Sampling continent | Host taxa | RdRp length | RdRp partial/complete | Catalytic triad | Top BLASTx hit | % identity |
|---|---|---|---|---|---|---|---|---|
| GBHO01039923.1 | Plant bug-associated ormycovirus 1 | North America | Hexapoda | 3277 | partial | NDD | Downy mildew lesion associated ormycovirus 6 | 41.0 |
| GBNA01013951.1 | Wasp-associated ormycovirus 1 | Africa | Hexapoda | 3056 | complete | NDD | Phytophthora cinnamomi ormycovirus 6-4 | 30.7 |
| GBTA01002218.1 | Moth-associated ormycovirus 2 | Oceania | Hexapoda | 801 | partial | NDD | Downy mildew lesion associated ormycovirus 6 | 38.8 |
| GBTA01054149.1 | Moth-associated ormycovirus 1 | Oceania | Hexapoda | 3801 | partial | NDD | Downy mildew lesion associated ormycovirus 6 | 34.5 |
| GDXN01051862.1 | Grasshopper-associated ormycovirus 1 | Europe | Hexapoda | 1475 | partial | NDD | Downy mildew lesion associated ormycovirus 6 | 35.1 |
| GEYJ01092710.1 | Mite-associated ormycovirus 1 | Europe | Chelicerata | 2589 | partial | NDD | Phytophthora cinnamomi ormycovirus 6-4 | 32.3 |
| GFJG01059483.1 | Crab-associated ormycovirus 1 | North America | Crustacea | 3396 | partial | NDD | Downy mildew lesion associated ormycovirus 6 | 40.0 |
| GHZM01086934.1 | Termite-associated ormycovirus 6 | Oceania | Hexapoda | 1223 | partial | NDD | Erysiphe lesion-associated ormycovirus 1 | 35.5 |
| GHZM01115244.1 | Termite-associated ormycovirus 7 | Oceania | Hexapoda | 2814 | partial | NDD | Phytophthora cinnamomi ormycovirus 11-3 | 29.0 |
| GHZM01131186.1 | Termite-associated ormycovirus 5 | Oceania | Hexapoda | 2977 | partial | NDD | Phytophthora cinnamomi ormycovirus 7-5 | 27.4 |
| GHZM01206470.1 | Termite-associated ormycovirus 4 | Oceania | Hexapoda | 2785 | partial | NDD | Downy mildew lesion associated ormycovirus 6 | 34.4 |
| GHZM01445935.1 | Termite-associated ormycovirus 2 | Oceania | Hexapoda | 2756 | partial | NDD | Botourmiaviridae sp. (UYL94578) | 32.0 |
| GIAM01545236.1 | Termite-associated ormycovirus 1 | Oceania | Hexapoda | 2903 | partial | NDD | Erysiphe lesion-associated ormycovirus 1 | 27.1 |
| GKFO01000268.1 | Beetle-associated ormycovirus 1 | Europe | Hexapoda | 1082 | partial | GDD | Downy mildew lesion associated ormycovirus 1 | 28.2 |
| HBDP01107766.1 | Bristletail-associated ormycovirus 1 | unknown | Hexapoda | 3120 | complete | GDD | Erysiphe lesion-associated ormycovirus 1 | 29.0 |
